# Deep Learning Prediction of Intact *N*-/*O*-Glycopeptide Tandem Mass Spectra Enhances Glycoproteomics

**DOI:** 10.1101/2025.07.20.665808

**Authors:** Yu Zong, Yuxin Wang, Liang Qiao

**Affiliations:** Department of Chemistry, and Minhang Hospital, Fudan University, Shanghai, China; Department of Computer Science, and Institute of Modern Languages and Linguistics, Fudan University, Shanghai, China

**Keywords:** *O*-glycosylation, glycoproteomics, deep-learning, graph neural network, mass spectrometry

## Abstract

Protein glycosylation, a post-translational modification involving the attachment of glycans to proteins, plays critical roles in numerous physiological and pathological cellular functions. Characterization of protein glycosylation is among one of the most challenging problems in proteomics due to the high heterogeneity of glycosites and glycan structures. Recently, deep learning has been adopted to predict *N*-glycopeptide tandem mass (MS/MS) spectra and exhibited a promising effect in *N*-glycoproteomics. However, current deep learning frameworks struggle to accurately predict *O*-glycopeptide MS/MS spectra due to the complexity of *O*-glycopeptides and the limited availability of training data. In this study, we introduce DeepGPO, a deep learning framework for the prediction of both *N*- and *O*-glycopeptide MS/MS spectra. The DeepGPO incorporates a Transformer module alongside two graph neural network (GNN) modules specifically designed for handling branched glycans. To address the issue of data scarcity in *O*-glycoproteomics datasets, various training methods are adopted in DeepGPO, such as the introduction of training weights for different MS/MS spectra and the adoption of pre-training strategies. DeepGPO exhibits accurate prediction of *N*- and *O*-glycopeptides MS/MS spectra. With the predicted MS/MS, glycosylation sites can be localized even in the absence of site-determining ions, for instance using higher-energy collisional dissociation (HCD) for the localization of *O*-glycosylation. We also explored the possibility of differentiating *O*-glycosites and *N*-glycosites using the predicted MS/MS spectra. DeepGPO primarily addresses mono-glycosylated peptides and is capable of handling doubly glycosylated peptides, but it currently cannot process peptides with three or more glycan modifications due to limited training data and increased spectral complexity. We anticipate that DeepGPO will inspire future advancements in glycoproteomics research.

## Introduction

Glycosylation is one of the most significant post-translational modifications (PTMs) of proteins, playing a crucial role in numerous physiological and pathological cellular functions^1^. In-depth research into glycosylation is critical in medical treatment, diagnosis and prognosis^2^. In protein glycosylation, glycans are usually attached to the side chains of asparagine (*N*-glycosylation) or serine/threonine (*O*-glycosylation). Glycosylation is highly complex, characterized by dynamically modified amino acids, varied monosaccharides composition, and intricate glycan structures. Despite the complexity of the branched structures of glycans for both *N*- and *O*-glycosylation, *O*-glycosylation presents even more significant analytical challenges because there is no specific *O*-glycosylation motif while any serine/threonine can be modified^3^. These heterogeneities pose substantial challenges in distinguishing glycopeptide isomers^4, 5^.

To date, liquid chromatography coupled tandem mass spectrometry (LC-MS/MS) is the leading method in glycoproteomics research. Tandem MS is used to characterize glycopeptides based on accurate molecular weight and specific fragment ions. One major challenge in glycoproteomics, particularly in *O*-glycoproteomics, is glycosite localization. Sequence database search engines, such as pGlyco3^6^, MSFragger-Glyco^7^ and MetaMorpheus^8^, typically rely on site-determining ions for glycosite localization, using MS/MS spectra generated from electron transfer dissociation (ETD), which can preserve glycan modifications during fragmentation. However, the efficiency of electron-based dissociation depends heavily on precursor ion charge density and often fails to produce sufficient fragments. To overcome this problem, ETD is frequently combined with higher-energy collisional dissociation (HCD) to generate more fragment ions, which nevertheless increases duty cycle times and reduces throughput. Alternatively, *O*-glycoproteases, such as OgpA^9, 10^, IMPa^11^ and StcE^12^, that can cleave proteins at sites adjacent to the appended glycans, are employed for *O*-glycosylation site identification. However, the specificity and cleavage efficiency of many enzymes remain suboptimal^3^.

Unlike sequence searching based methods, spectra-match is used for peptide identification by matching experimental mass spectra with a spectral library, utilizing both intensity and m/z information. This approach overcomes the need for the observation of site-determining fragment ions for glycosite localization and improves identification sensitivity^13^. However, constructing glycopeptide MS/MS spectral libraries remains challenging due to the difficulty of glycopeptide synthesis and the complexity of biological samples. As an alternative, deep-learning prediction of peptide MS/MS spectra offers an efficient method for constructing spectral libraries. Various deep learning-based MS/MS spectra prediction tools, such as pDeep series^14–16^, Prosit^17^, DeepMass:Prism^18^, DeepDIA^19^, DeepPhospho^20^, DeepFLR^21^, DeepGlyco^22^ and DeepGP^23^, have been developed. DeepGlyco^22^ and DeepGP^23^ are recently published deep-learning models for *N*-glycopeptides MS/MS spectra prediction. Despite these advancements, none of these tools can predict MS/MS spectra for *O*-glycopeptides due to the complexity of *O*-glycopeptides and the scarce training data.

Herein, we present a deep-learning based framework, DeepGPO, for *N*-/*O*-glycopeptides MS/MS spectra prediction and *O*-glycosite localization. DeepGPO is composed of a Transformer module alongside two graph neural network (GNN) modules specifically designed for handling branched glycans. Advanced training techniques are employed to mitigate the limited availability of training data. DeepGPO has explored the potential of spectral searching for *O*-glycopeptides and realized *O*-glycosite localization based on HCD MS/MS spectra. Besides, DeepGPO shows promising results in distinguishing glycopeptides as *N*-glycopeptides or *O*-glycopeptides.

## Results

### DeepGPO framework

DeepGPO is developed based on our recently published model DeepGP^23^ for *N*-glycopeptides. The deep learning model of DeepGPO incorporates a Transformer module alongside two graph neural network (GNN) modules for handling branched glycans, similar to DeepGP (**Supplementary Fig. 1**). DeepGPO is capable of predicting *N*- and *O*-glycopeptides MS/MS spectra (**Fig. 1a**). B/Y and b/y ions are considered concurrently to preserve the relative intensity of the fragment ions. Up to 16 types of B/Y fragment ions or 24 types of b/y fragment ions are considered for each cleavage event by combining various neutral losses and charge states, as detailed in the **Methods** section. To improve DeepGPO’s generalization across *N*- and *O*-glycopeptides, no restrictions are imposed on glycosites or glycan types. Besides, DeepGPO considers the common modifications, including Oxidation [M], Acetyl [Protein N-term] and Carbamidomethyl [C], and two modifications often encountered in glycoproteomics, namely Deamidated [N] and Guanidinyl [K]. Deamidation occurs when PNGase F is used to remove *N*-glycosylation, and guanidination of lysine is frequently applied to prevent conjugation of lysine residues to aldehyde groups on solid phase and to enable the identification of glycopeptides containing lysine residues^10^. DeepGPO supports any possible glycopeptide length and precursor charge state. These features make DeepGPO a highly versatile model for glycopeptides analysis.

**Figure 1.**
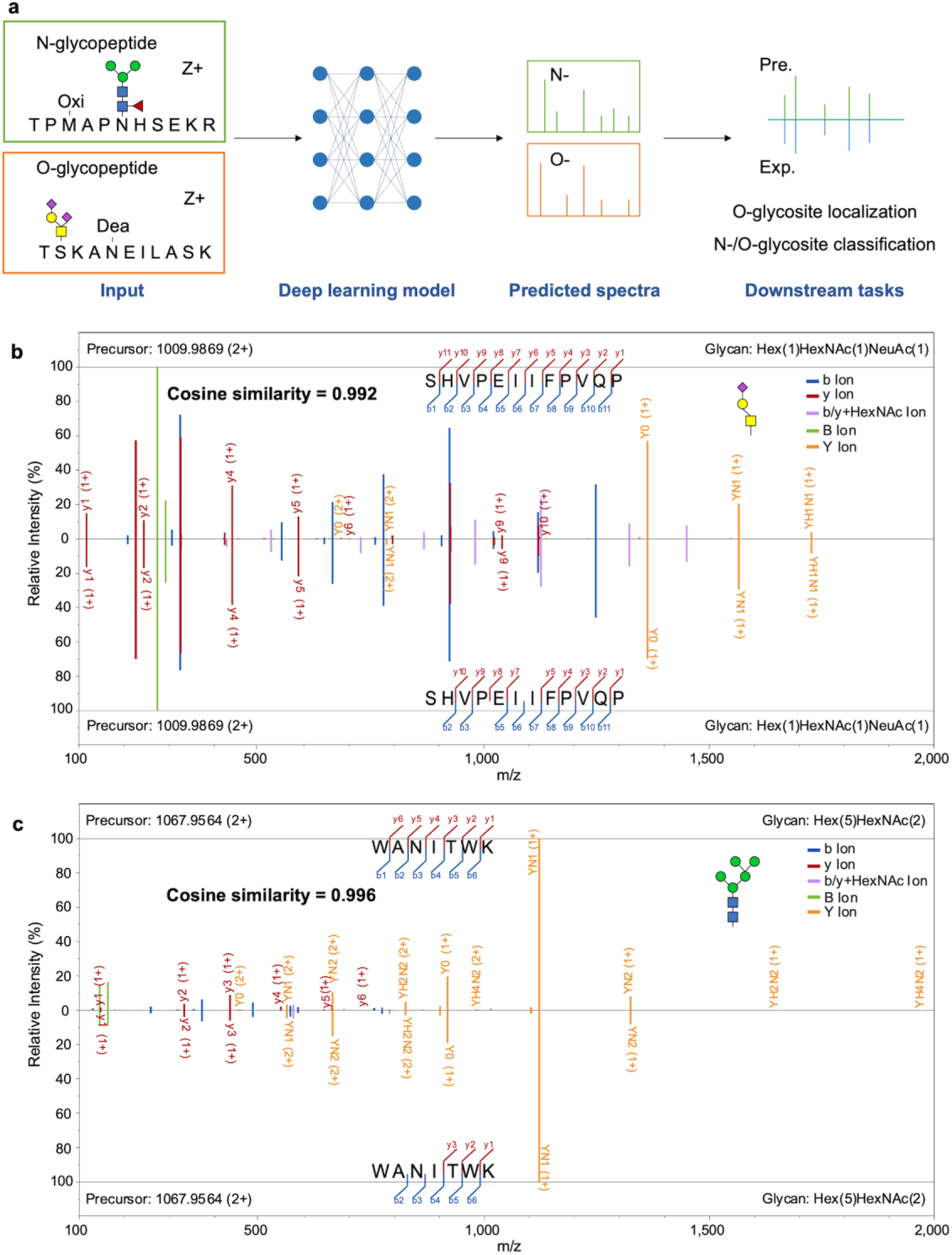
DeepGPO framework and *N*-/*O*-glycopeptide MS/MS spectra prediction. **(a)** The framework of DeepGPO. Comparison of the experimental MS/MS with the predicted MS/MS of **(b)** an *O*-glycopeptide and (**c)** an *N*-glycopeptide. Top: the predicted MS/MS spectra; Bottom: the experimental MS/MS spectra. Experimental MS/MS spectra were pre-processed to remove noise peaks for clearer comparison.

In DeepGPO, HCD MS/MS spectra serve as the training, validation and test data. For HCD-pd-EThCD, DeepGPO utilizes the parent HCD MS/MS spectra, while glycopeptide identification is performed by pGlyco3 using the parent HCD MS/MS spectra and the triggered EThCD MS/MS spectra (**Methods**). DeepGPO adopts sqrt-cosine similarity (**Methods**) as the primary metric for glycopeptide MS/MS prediction, with the consideration that peak intensities follow a Poisson distribution and that the square root transformation can stabilize the peak intensity variance^24^. This transformation is particularly advantageous as it assigns greater weight to low-intensity peaks, thereby enhancing the prediction accuracy for low-abundance ions. Significantly, DeepGPO incorporates any peak identified as a fragment of glycopeptides into its metric calculation, not just those predicted by DeepGPO. Beyond sqrt-cosine, DeepGPO also employs additional metrics for performance representation, including cosine similarity (COS) and Pearson correlation coefficient (PCC).

Given the substantial scarcity of *O*-glycoproteomics training data, we implement three strategies to optimize DeepGPO’s performance. First, we retain all glycopeptide MS/MS spectra during model training without removing duplicates. Critically, we ensure no glycopeptides overlap between the training and validation/test datasets. This approach acts as a form of data augmentation, where the intrinsic variations in different MS/MS spectra for the same glycopeptide replace manual augmentation methods. Second, we introduce a loss re-weighting method. **Supplementary Fig. 2** illustrates the workflow for assigning training weights to glycopeptide MS/MS spectra by considering the pGlyco3 identification results and the glycoprotease specificity where available. A higher weight indicates a higher confidence in the identification result. During model training, only MS/MS spectra with weight exceeding 0.5 are considered, and these weights are integrated into the loss function by multiplying the loss function by the corresponding weight for each MS/MS spectrum. As a result, inaccurate predictions for MS/MS spectra with higher weights incur a greater penalty for DeepGPO. This approach preserves a wider range of training MS/MS spectra while allowing the model to quantitatively learn from more reliable MS/MS spectra, avoiding the need for stringent filtering rules that would significantly reduce the pool of available *O*-glycopeptides MS/MS spectra.

Furthermore, a pre-training strategy is implemented. We have trained DeepGPO based on Transformer without pre-training^25^, BERT^26^ and DeepGP^23^. These models are of the same model configuration, but varied on the usage of pre-training. Transformer starts with randomly initialized parameters. BERT benefits from pre-training with unlabeled natural language. DeepGP is a BERT-based framework for *N*-glycopeptides MS/MS spectra prediction. The benchmarking dataset (Dataset 1) comprises 18,260 *O*-glycopeptides MS/MS spectra with weight exceeding 0.5 (PXD037415, **Supplementary Table 1**, **Supplementary Note 1**), previously published by Suttapitugsakulet al.^27^. The dataset was divided into the training and the validation dataset at a 9:1 ratio. The model’s performance was evaluated on the validation dataset during training using different base models. The result showed that DeepGPO based on DeepGP can reach the highest median sqrt-cosine in the prediction of *O*-glycopeptide MS/MS spectra while requiring the least epochs (**Supplementary Fig. 3**), indicating that the pre-training strategy could benefit downstream tasks with limited data. In the following experiments, only DeepGP was considered as the base model. Multi-stage fine-tuning could further enhance the performance. Based on the model trained using other *O*-glycoproteomics datasets (Dataset 2-11^9, 28–35^, **Supplementary Table 1**), DeepGPO can achieve better performance on the same validation data from Dataset 1 (**Supplementary Fig. 4**).

**Fig. 1b** and **Fig. 1c** show the prediction performance using one *N*-glycopeptide and one *O*-glycopeptide as examples. The two glycopeptides are both from the Dataset 1 but with different experimental conditions as described in **Supplementary Note 1**. The *N*-glycopeptide spectrum is acquired by sceHCD and the *O*-glycopeptide spectrum is the parent HCD MS/MS spectrum from the HCD-pd-EThCD MS/MS spectrum. Due to the different experimental conditions of *N*- and *O*-glycopeptides MS/MS spectra, DeepGPO was trained separately. The model for the prediction of the *O*-glycopeptide MS/MS spectra was the Finetuned-DeepGPO model in **Supplementary Fig. 4**. For the prediction of the *N*-glycopeptide MS/MS spectra, DeepGPO was trained with 18,836 *N*-glycopeptide MS/MS spectra of Dataset 1. The results demonstrate that the predicted MS/MS spectra for both glycopeptides closely resemble their corresponding experimental MS/MS spectra (**Fig. 1b, c**). DeepGPO successfully captured the fragmentation pattern of both *N*-glycopeptides and *O*-glycopeptides.

### Performance evaluation for *O*-glycopeptide MS/MS prediction by DeepGPO

The performance of glycopeptide MS/MS prediction by DeepGPO was benchmarked using Dataset 1^27^ (**Supplementary Table 1**). Dataset 1 employed IMPa that cleaves the N-terminal of a serine or threonine residue containing an *O*-glycan modification. The single-shot experiments on mouse brain tissues in Dataset 1 served as the test data, while all the other *O*-glycopeptides data in Dataset 1 served as the training data (**Supplementary Note 1**). The glycan types of the test data are shown in **Supplementary Fig. 5**. Glycopeptides presented in the training data were intentionally removed from the test data to ensure representation accuracy. We observed a median cosine similarity of 0.931 for *O*-glycopeptide MS/MS prediction (**Fig. 2a** *Model1*). Moreover, by using model pre-trained with other *O*-glycoproteomics datasets (Dataset 2-11^9, 28–35^, **Supplementary Table 1**) and then fine-tunned with the training data of Dataset 1, the median cosine similarity can be further improved to 0.952 (**Fig. 2a** *Model2*). We conducted further analysis on the results of **Fig. 2a** *Model2*. Upon narrowing our focus to the glycan B/Y ions, the accurate prediction of ion intensity was also confirmed, with the median cosine similarity of 0.983 (**Fig. 2a** *BY*). Beyond cosine similarity, we further explored the distribution of Pearson correlation coefficients (PCC) between the predicted and experimental MS/MS spectra (**Fig. 2b**) with the same setting as **Fig. 2a** *Model2*, and a high median PCC of 0.95 was also demonstrated.

**Figure 2.**
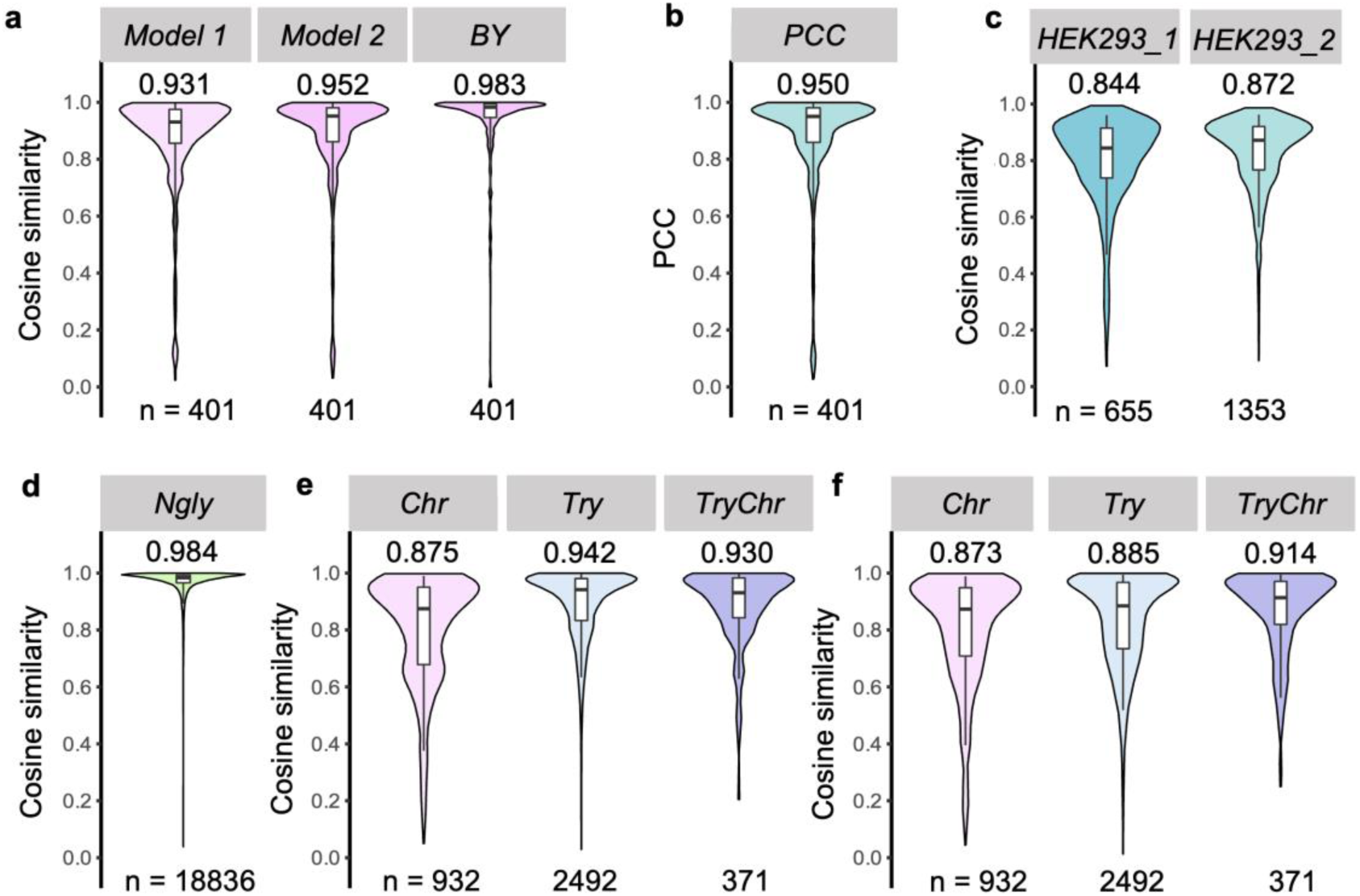
Performance of DeepGPO in MS/MS prediction. **(a)** The distribution of cosine similarity between predicted and experimental MS/MS spectra of *O*-glycopeptides from Dataset 1 mouse brain samples. Model1: model without pretraining. Model2: model with pretraining by other *O*-glycoproteomics datasets. BY: consideration of only B/Y glycan ions for cosine similarity. **(b)** The distribution of pearson correlation coefficient computed between the predicted and experimental MS/MS spectra of *O*-glycopeptides from Dataset 1 mouse brain samples. The DeepGPO model is the Model2. (**c**) The distribution of cosine similarity between the predicted and experimental MS/MS spectra of *O*-glycopeptides from Dataset 1 HEK293 cells. HEK293_1 represents the parent HCD MS/MS spectra from HCD-pd-EThCD and HEK293_2 represents the HCD MS/MS spectra without triggered EThCD. **(d)** The distribution of cosine similarity between the predicted and experimental MS/MS spectra for the *N*-glycopeptides from Dataset 1. (**e**) The distribution of cosine similarity between the predicted and experimental MS/MS spectra for the glycopeptides digested with trypsin (Try), chymotrypsin (Chr) and the combination of trypsin and chymotrypsin (TryChr) from Dataset 3. The DeepGPO was pre-trained by other *O*-glycoproteomics datasets, and the fined tunned on the data split from Dataset 3. (**f**) The distribution of cosine similarity between the predicted and experimental MS/MS spectra for Dataset 3. No pre-training was applied. The medians are indicated. The boxes and whiskers show the quantiles and 95% percentiles, respectively. The numbers of spectra for test are indicated below each graph. Source data are provided as a Source Data file.

In another analysis, the *O*-glycopeptides from HEK293 cells in Dataset 1 were used as the test data, while the *O*-glycopeptides from the mouse brain samples in Dataset 1 served as the training data. The DeepGPO model was trained firstly using other *O*-glycoproteomics datasets (Dataset 2-11^9, 28–35^, **Supplementary Table 1**), and then finetuned by the training data of Dataset 1. For the parent HCD MS/MS spectra from HCD-pd-EThCD and the HCD MS/MS spectra (without triggered EThCD) (**Supplementary Note 1**), the cosine similarity reached 0.844 and 0.872, respectively (**Fig. 2c**). Given that the organism of the training data was different from the test data, the cosine similarity was decreased compared to those in **Fig. 2a**, but was not bad, demonstrating that DeepGPO can be applied to multiple organisms and has cross-organisms applicability.

For Dataset 1, the same mouse brain tissues were used for both *N*- and *O*-glycoproteomics analyses by different enrichment methods and experimental conditions (**Supplementary Note 1**). Due to the significant differences in experimental conditions, we used DeepGPO trained by the *N*-glycopeptides data to predict the MS/MS spectra of *N*-glycopeptides. The results showed a median cosine similarity of 0.984 (**Fig. 2d**), demonstrating that DeepGPO can accurately predict both *O*- and *N*-glycopeptide MS/MS spectra. It is noted that the performance of the deep-learning model on *N*-glycopeptides is consistent with that of DeepGP^23^, and no further discussion on the prediction of *N*-glycopeptides MS/MS spectra is made here.

In addition to Dataset 1, we also benchmarked DeepGPO on Dataset 3^29^ (PXD004590, **Supplementary Table 1**, **Supplementary Note 1**), which involved human samples digested with trypsin (Try), chymotrypsin (Chr) and the combination of trypsin and chymotrypsin (TryChr). Each condition served as the test data in turn, with the remaining for training. DeepGPO was firstly trained with other *O*-glycoproteomics datasets (Dataset 1-2, 4-11^9, 27, 28, 30–35^, **Supplementary Table 1**) before being fine-tuned on the data split from Dataset 3. DeepGPO demonstrated remarkable accuracy in predicting glycopeptide MS/MS spectra (**Fig. 2e**). When comparing results without pre-training (**Fig. 2f**), the performance of *O*-glycopeptides MS/MS prediction on the Try data showed the most significant improvement, while that on the Chr dataset remained consistent. This is attributed to the absence of chymotrypsin-digested peptides in the pre-training data from Dataset 1-2, 4-11. For Dataset 3, no *O*-glycoprotease was used, demonstrating that DeepGPO can be applied to *O*-glycopeptides with glycosylation at any positions.

### *O*-glycosite localization from HCD MS/MS spectra by DeepGPO

With the accurate prediction of glycopeptides MS/MS spectra, DeepGPO can be used for *O*-glycosite localization. The principle was first illustrated through representative examples. Two HCD MS/MS spectra from Dataset 2^28^ (PXD018560, **Supplementary Table 1**, **Supplementary Note 1**) sharing the same peptide sequence and glycan modification but different glycosites (site 1 or 6) were collected. The sites were both verified by the corresponding ETD MS/MS spectra (**Supplementary Fig. 6**). *In silico* HCD MS/MS spectra were generated for the two glycopeptides by DeepGPO. As shown in **Fig. 3** and **Supplementary Fig. 7**, when the glycosylation sites were consistent between the predicted HCD MS/MS spectrum and the experimental one, a significant higher spectrum similarity was observed, i.e. sqrt-cosine similarity of 0.978 for both sites as 6 and 0.975 for both sites as 1; while the similarity was lower when the two sites were different, i.e. 0.873 for predicted site at 1 against experimental site at 6 and 0.849 for predicted site at 6 against experimental site at 1. Further examples are shown in **Supplementary Fig. 8**.

**Figure 3.**
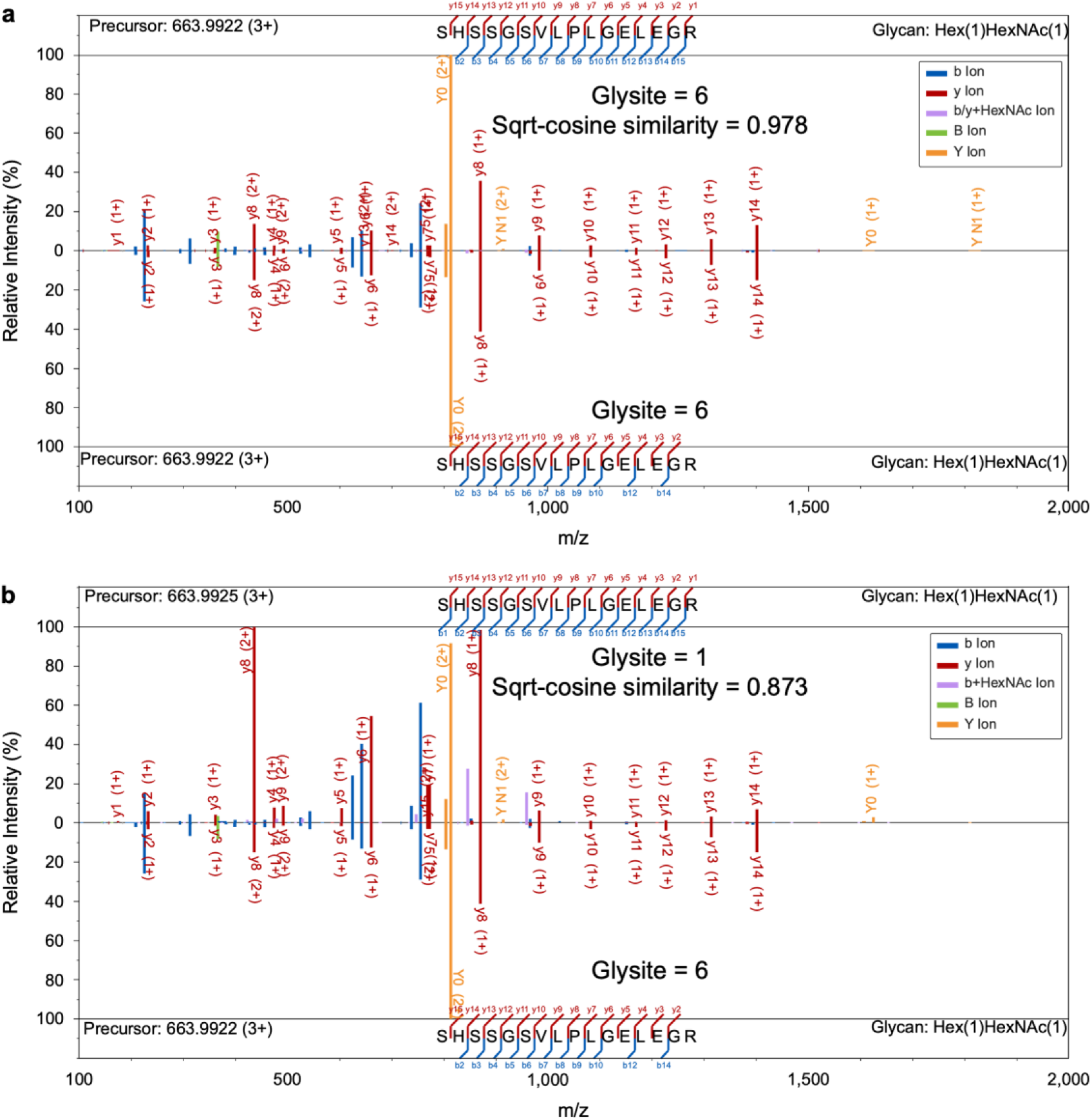
Comparison of experimental and predicted HCD MS/MS of glycopeptides with the same peptide sequence and glycan modification but different glycosites. (a) Glycosite = 6 for both predicted and experimental MS/MS spectra; (b) Glycosite = 1 for predicted MS/MS spectra and Glycosite = 6 for experimental MS/MS spectra. Top: the predicted MS/MS spectra; Bottom: the experimental MS/MS spectra. Experimental spectra were pre-processed to remove noise peaks for clearer comparison.

Then, the DeepGPO was applied to *O*-glycosite localization on large-scale datasets. We have firstly evaluated the possibility using Dataset 1. The test data were the single-shot experiments on the mouse brain tissues. The DeepGPO model was trained using the Dataset 1. Glycosite localization was performed on the MS/MS spectra identified as mono-*O*-glycosylated peptides with multiple candidate glycosites and starting with Ser/Thr. The spectra with ambiguous glycosite localization were excluded, where pGlyco3 reported site-groups containing multiple indistinguishable sites. The candidate glycopeptides list was generated from the pGlyco3 search result considering all possible glycosites. sqrt-cosine similarity was calculated between the predicted and the experimental MS/MS. The top match was reported for each experimental MS/MS (**Fig. 4a**). As the sample was prepared by IMPa *O*-glycoprotease, we considered the correct glycosites as N-termini. With the parent HCD MS/MS spectra of the HCD-pd-EThCD, DeepGPO localized 65 MS/MS spectra with glycosites different from N-termini out of the 1723 tested MS/MS spectra, demonstrating an empirical false localization rate (FLR) of 3.8%. In contrast, pGlyco3, using EThCD, localized 116 MS/MS spectra with glycosites different from N-termini. At a 5% empirical FLR, DeepGPO retained all the 1723 MS/MS spectra, while pGlyco3 localized 1495 (**Fig. 4b**, **Supplementary Data 1**). During pGlyco3 analysis, no cutoff on localization probability was applied.

**Figure 4.**
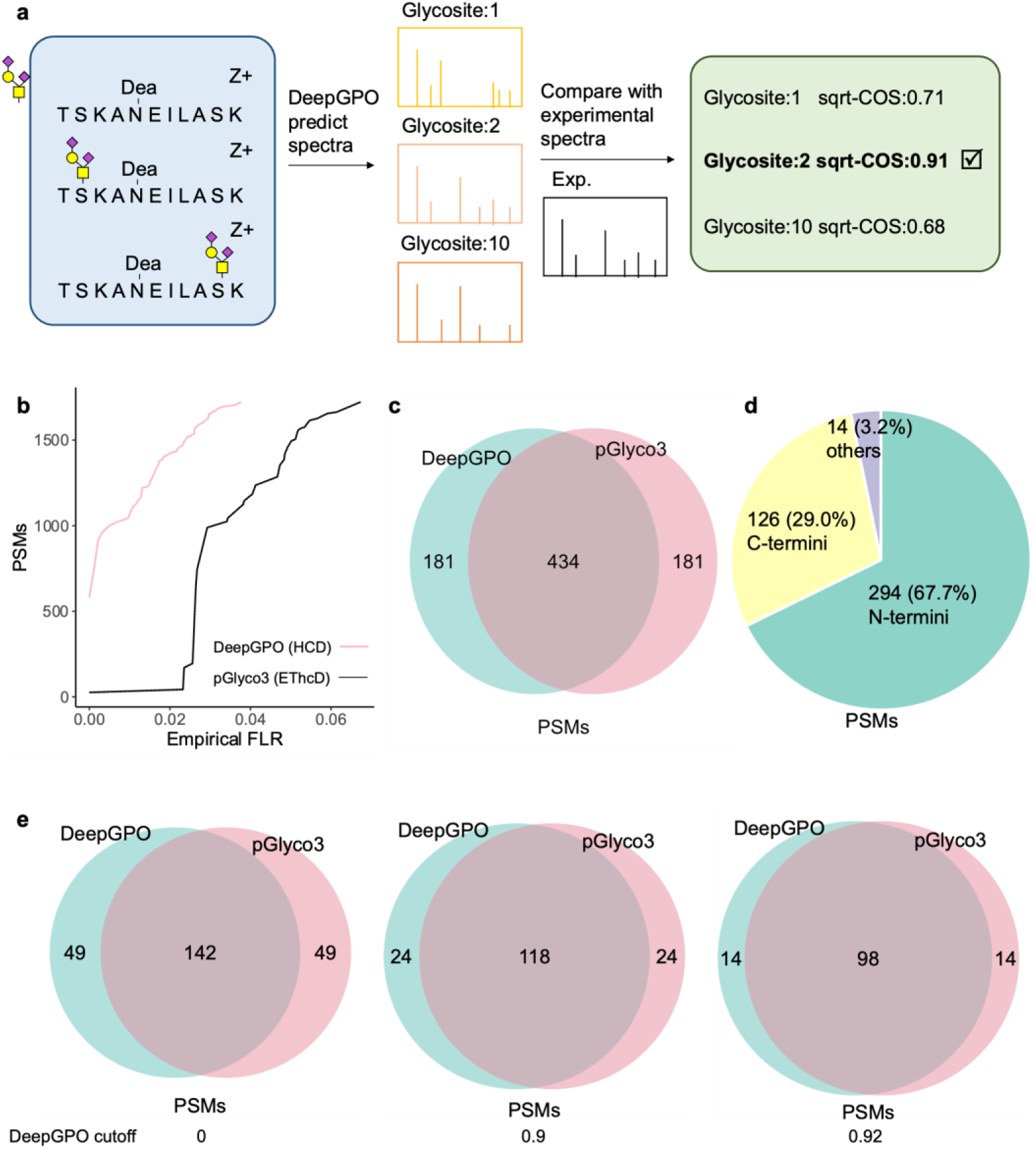
Performance of DeepGPO on glycosite localization. **(a)** Protocol of performing glycosite localization by DeepGPO. **(b)** Comparison of the number of identified glycopeptide PSMs under a given empirical FLR by DeepGPO from HCD and by pGlyco3 from HCD-pd-EThCD for Dataset 1. **(c)** Venn diagram of the number of PSMs identified by DeepGPO and pGlyco3 for Dataset 4. pGlyco3 analyzed the EThcD while the DeepGPO analyzed the HCD MS/MS spectra from the EThcD. (**d**) Pie chart of the PSMs shared by DeepGPO and pGlyco3 with N-termini glycosylation, C-termini glycosylation and other position glycosylation for Dataset 4. (**e**) Venn diagram of the number of PSMs identified by DeepGPO with a similarity score cutoff of 0, 0.9 and 0.92, and by pGlyco3 for Dataset 3. The pGlyco3 glycosite localization probability was ≥ 0.8. pGlyco3 analyzed the EThcD while the DeepGPO analyzed the HCD spectra from the EThcD. Source data are provided as a Source Data file.

In addition to Dataset 1, DeepGPO was also benchmarked on Dataset 4, where digestions were performed with AM0627 or AM0627 mutants (AM0627^W149A^, AM0627^F290A^, AM0627^Y287A^) against recombinant glycoproteins podocalyxin, MUC16, PSGL-1, and CD43^34^. The glycopeptides were analyzed by HCD-pd-EThCD. The test data were from the glycopeptides digested with AM0627, which cleaved between adjacent residues carrying truncated core 1 *O*-glycans. As a result, it is hypothesized that the glycosite was located at either the N- or C-termini of the peptides. The spectra with ambiguous glycosite localization were excluded. The DeepGPO model was trained using the Dataset 4. From the test data of Dataset 4 (**Supplementary Note 1**), we obtained 615 MS/MS spectra corresponding to 3202 candidate *O*-glycopeptides. DeepGPO, relying solely on HCD spectra, localized 131 MS/MS spectra with glycosites different from either the N or C-termini of the peptides. In comparison, pGlyco3, using EThCD, localized 125 MS/MS spectra with glycosites different from either the N or C-termini of the peptides. During pGlyco3 analysis, no cutoff on localization probability was applied. The Venn diagram of PSMs identified by pGlyco3 and DeepGPO is shown in **Fig. 4c**. Among the 615 MS/MS spectra, 434 yielded identical results between pGlyco3 and DeepGPO, including 294 (294/434 = 67.7%) with the glycosite at the N-termini, 126 (126/434 = 29.0%) with the glycosite at the C-termini, and 14 (14/434 = 3.2%) with other glycosites (**Fig. 4d**). It is noted that although specific enzymatic cleavage should occur between adjacent residues carrying truncated core 1 *O*-glycans, there can be non-specific cleavages during enzymatic reaction. The observed *O*-glycopeptides identification results are consistent with the specificity of the enzyme.

In Dataset 1 and 4, specific *O*-glycoproteases were used, which may introduce bias to the glycosite localization performance evaluation. To demonstrate a more generalized application, we further benchmarked the method on Dataset 3, which consists of HCD-pd-EThCD MS/MS spectra of glycopeptides generated with different proteases, including trypsin, chymotrypsin and the combination of trypsin and chymotrypsin, but without *O*-glycoprotease. The data with the combination of trypsin and chymotrypsin were used as the test data; while the others in the dataset were used as the training data. The glycan sites were determined by pGlyco3 based on the EThCD MS/MS spectra with a localization probability ≥ 0.8. The spectra with ambiguous glycosite localization were excluded. In total, 191 HCD MS/MS spectra were collected corresponding to 622 candidate *O*-glycopeptides. The identification results by DeepGPO from HCD MS/MS spectra were largely consistent (142/191 = 74.3%) with the identification results by pGlyco3 from the EThCD MS/MS spectra (**Fig. 4e**). We also collected the HCD MS/MS spectra of glycopeptides identified by pGlyco3 with localization probability < 0.8. As shown in **Supplementary Fig. 9**, for the low-confidence group (83 spectra), only 51 showed consistent localization, resulting in a lower consistency rate of 61.4%. All the benchmarking results demonstrate that DeepGPO can perform glycosites localization using only HCD MS/MS spectra.

It is noted that the top-match by spectra similarity is not necessarily correct. To further improve the identification accuracy by DeepGPO, we applied a cut-off on the spectra similarity score (sqrt-cosine similarity). **Supplementary Fig. 10** shows the score distributions for both correct and incorrect PSMs across the three datasets (Datasets 1, 3, and 4). For Dataset 1, the correct PSMs are those identified with glycosylation on the N-terminus. For Dataset 4, the correct PSMs are those identified with glycosylation on the N- or the C-terminus. For Dataset 3, the correct PSMs are those consistent with the pGlyco3 identification result using EThcD (with glycosite localization probability ≥ 0.8). As shown in **Supplementary Fig. 10**, the distributions of correct and incorrect PSMs are separated at the sqrt-cosine similarity around 0.90. Based on this, we applied the similarity score thresholds of 0.90 and 0.92 to the Dataset 3 identification result by DeepGPO, and further compared to the pGlyco3 identification result (**Fig. 4e**). At the threshold of 0.9, the consistency between DeepGPO and pGlyco3 was 83.1% (118/142). At the threshold of 0.92, the consistency between DeepGPO and pGlyco3 was 87.5% (98/112). On glycosite level, the consistency between DeepGPO and pGlyco3 were 65.8%, 72.5% and 76.7% for sqrt-cosine similarity threshold of 0, 0.90 and 0.92, respective (**Supplementary Fig. 11)**. The consistency at the protein glycosites level was lower compared to the PSMs level, which also indicated that the protein glycosites identified by multiple MS/MS spectra were more confident. Considering that the pGlyco3 identification result can also contain false results, such consistency between DeepGPO and pGlyco3 is reasonable. It is noted that the pGlyco3 glycosite identification result is based on ETD, while the DeepGPO localizes glycosites solely based on HCD.

Glycosylation plays a crucial role in infection and immune evasion. Dataset 5^30^ (PXD022896, **Supplementary Table 1**, **Supplementary Note 1**) reports *O*-glycosylation sites for insect cell-expressed S protein and human cell-expressed S protein S1 subunit of SARS-CoV-2 using EThCD and HCD. We evaluated our method for *O*-glycosite localization using Dataset 5 with an emphasis on the human cell-expressed S protein S1 subunit digested by trypsin. Due to the limited size of Dataset 5, it was not feasible to fine-tune DeepGPO on this dataset. Instead, a model trained with all the *O*-glycopeptide datasets (Dataset 1-11) was used. DeepGPO identified 15 glycosylation sites (22, 33, 114, 286, 316, 323, 325, 547, 638, 645, 659, 673, 676, 678, 680) based on the HCD MS/MS spectra (**Supplementary Data 2**). In the original article^30^, 14 glycosylation sites were reported based on EThCD for the human cell-expressed S protein S1 subunit, including 9 sites (286, 323, 325, 638, 659, 673, 676, 678 and 680) reported by our method.

### Analysis of doubly *O*-glycosylated peptides by DeepGPO

*O*-glycoproteomics features challenges on multiple-glycosylation due to the complexity of assigning glycan compositions to multiple glycosites. Here, we extended the application of DeepGPO to doubly *O*-glycosylated peptides, utilizing a publicly available dataset (Dataset 12, PXD017646, **Supplementary Table 1**, **Supplementary Note 1**) featuring multiply *O-*glycosylated peptides^36^. HCD-pd-ETD and HCD-pd-EThcD MS/MS spectra from the dataset for glycopeptides containing no more than two glycosylation modifications were selected for analysis. The MS/MS spectra was partitioned by unique glycopeptides at a 9:1 ratio, resulting in 976 spectra for training and 105 for test, with no glycopeptide overlap between the training and test sets. DeepGPO was first trained by using other *O*-glycoproteomics datasets (Dataset 1-11) before being fine-tuned on the training data of Dataset 12. Identification results reported in the original publication were used to label the HCD MS/MS spectra. On the entire test set, DeepGPO achieved a median cosine similarity of 0.974, demonstrating high alignment in HCD MS/MS prediction (**Fig. 5a**). Notably, for the subset of 70 doubly glycosylated peptides in the test set, the model maintained a high median cosine similarity of 0.970, indicating that the performance of HCD MS/MS prediction remained robust even for multiply glycosylated peptides. An illustrative example for one such peptide is provided in **Fig. 5b**, where the predicted and experimental MS/MS spectra show strong agreement.

**Figure 5.**
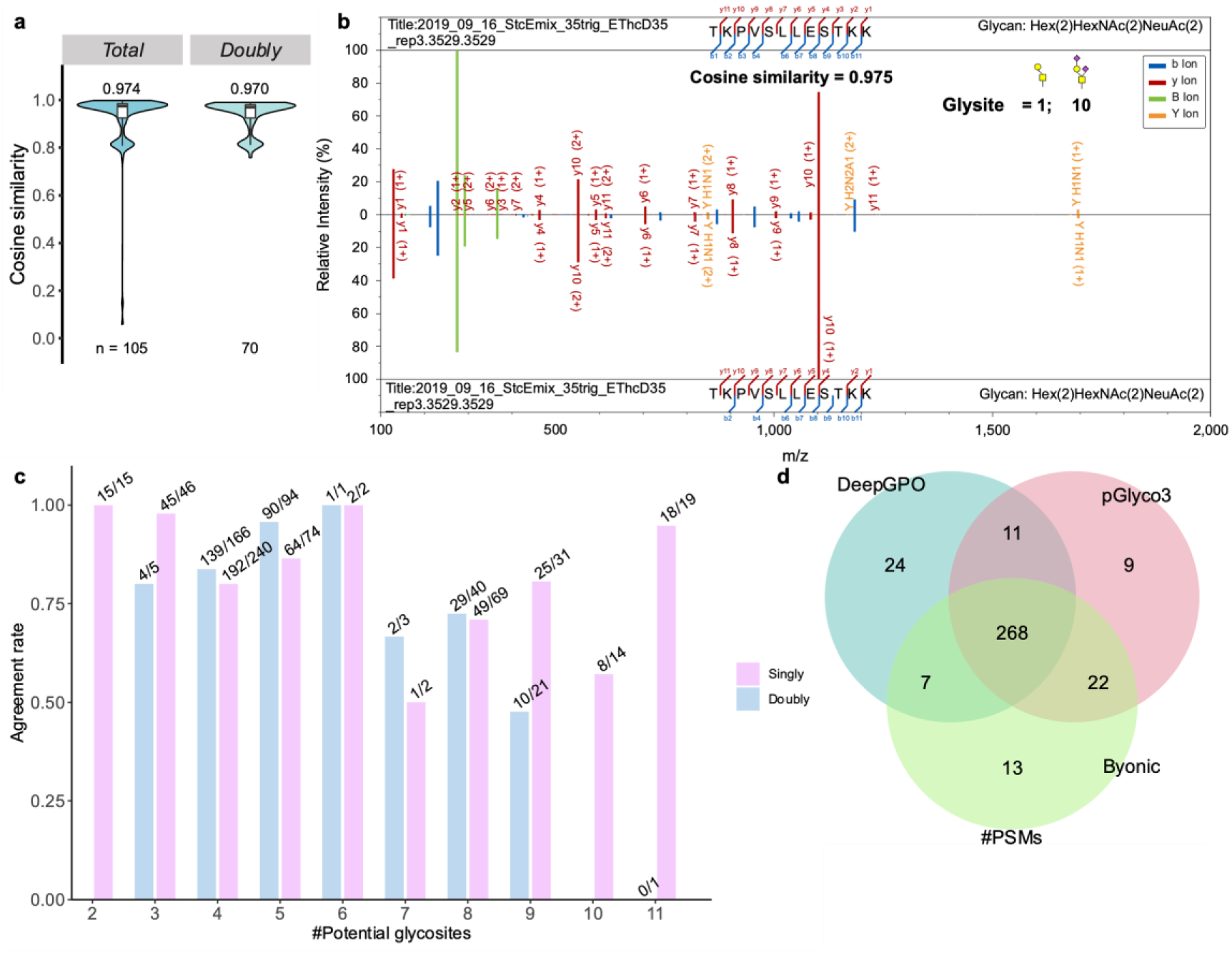
Performance of DeepGPO on multiply *O*-glycosylated peptides. (**a**) Cosine similarity between predicted and experimental HCD MS/MS spectra for *O-* glycopeptides from Dataset 12. Total: glycopeptides from the test dataset. Doubly: doubly *O-*glycosylated peptides split from the test dataset. The medians are indicated. The boxes and whiskers show the quantiles and 95% percentiles, respectively. The numbers of spectra for test are indicated below each graph. (**b**) Example of a doubly *O-* glycosylated peptide with glycosylation at positions 1 and 10. Top: predicted spectrum; bottom: experimental spectrum. Experimental MS/MS spectra were pre-processed to remove noise peaks for clearer comparison. (**c**) Localization consistency of DeepGPO compared to Byonic across peptides with different numbers of serine/threonine residues in Dataset 12. Bars show consistent spectra counts over total, separated by singly (blue) and doubly (pink) glycosylated peptides. (**d**) Venn diagram showing the overlap of glycopeptide-spectrum matches (PSMs) identified by pGlyco3, Byonic, and DeepGPO in Dataset 12. Source data are provided as a Source Data file.

We then assessed the localization accuracy in the case of doubly *O-*glycosylated peptides using DeepGPO. MS/MS spectra identified as glycopeptides with unambiguous glycosites were excluded (the number of glycans = the number of candidate sites). For singly glycosylated peptides, glycosylation site candidates were defined as all serine and threonine residues, consistent with previous analyses. For the doubly glycosylated peptides, one glycosite was fixed while the second glycan was allowed to localize across all serine and threonine residues. As shown in **Fig. 5c**, DeepGPO achieved high consistency with the original assignments for both singly and doubly glycosylated peptides. We further tested the MS/MS spectra only in the test dataset. Among 101 HCD MS/MS spectra, DeepGPO localized 77 cases in agreement with the reference.

To further strengthen the result, we also analyzed Dataset 12 using pGlyco3. A total of 310 MS/MS spectra were identified by both pGlyco3 and Byonic, with identical peptide sequences, charge states, and glycans. These MS/MS spectra were also consistently identified as either singly or doubly *O-*glycosylated peptides. As shown in **Fig. 5d**, among the 290 spectra for which the glycosylation sites were consistently assigned by pGlyco3 and Byonic, DeepGPO achieved the same localization in 268 cases, corresponding to a 92.4% agreement rate, highlighting the model’s competitive performance relative to established tools. Notably, among the 127 doubly *O-* glycosylated peptide MS/MS spectra within the 290 cases, DeepGPO’s results matched those of both pGlyco3 and Byonic for 117 spectra, yielding an agreement rate of 92.1% (117/127). It needs to emphasize that both pGlyco3 and Byonic employed ETD MS/MS spectra for glycosite localization, while DeepGPO used only HCD MS/MS spectra. It must be acknowledged that the method currently cannot handle peptides with three or more glycan modifications due to limited training data.

### Differentiation of *N*- and *O*-glycopeptides

As proteins usually undergo both *O*- and *N*-glycosylation, it is important to perform co-identification of *N*-/*O*-glycopeptides^37^. Current software solutions typically address the localization of *N*-glycopeptides and *O*-glycopeptides separately, wherein a single MS/MS spectrum might be identified as both a *N*-glycopeptide and an *O*-glycopeptide in separate rounds of database searches. To mitigate this issue, current methods often exclude glycopeptides containing the "NXS/T" sequon from being considered for *O*-glycosylation. While this approach reduces overlaps of MS/MS spectra in identification, it diminishes the sensitivity for *O*-glycopeptide detection. This limitation hinders the comprehensive understanding of protein glycosylation, highlighting the need for more refined methods to accurately detect and differentiate *N*- and *O*-glycosylation.

Mucins are a family of high molecular weight proteins produced by epithelial tissues, characterized by repetitive structures containing a high frequency of *N*- and *O*-glycosylation^38^. Dataset 13^38^ (PXD024995, **Supplementary Table 1**, **Supplementary Note 1**) contains HCD MS/MS spectra of mucin proteins. We utilized pGlyco3 to analyze the dataset in two separate rounds, operating in either *O*-glycan mode or *N*-glycan mode. pGlyco3 identified 7861 MS/MS spectra as *N*-glycopeptides and 5200 MS/MS spectra as *O*-glycopeptides. Among them, 223 MS/MS spectra were co-identified by both modes. DeepGPO was trained with the *N*-glycopeptides MS/MS spectra or the *O*-glycopeptides MS/MS spectra, excluding the 223 co-identified MS/MS spectra. The trained DeepGPO was subsequently tested on the co-identified MS/MS spectra. All the potential *O*-glycosites and *N*-glycosites were considered. MS/MS spectra were predicted for the *N*-glycopeptides and *O*-glycopeptides, and compared with the experimental MS/MS to report the one with the best spectral match, as illustrated in **Fig. 6a**. Among all the 223 MS/MS spectra, DeepGPO identified 196 as *O*-glycopeptides and 27 as *N*-glycopeptides (**Fig. 6b**, **Supplementary Data 3**). The glycan types of the identified glycopeptides are shown in **Fig. 6c**. For core-1 *O*-glycans, which lack GlcNAc, the 138/144 intensity ratio (m/z 138 to m/z 144) is expected to be small and close to 1^38^. As the glycopeptides identified from the 223 MS/MS spectra contain a large portion of core-1 *O*-glycans (**Fig. 6c**), it is expected that the 138/144 intensity ratio should be smaller for the *O*-glycopeptides than the *N*-glycopeptides, consistent with the observation (**Fig. 6d**, MS/MS spectra without the peak of m/z 144 excluded for comparison).

**Figure 6.**
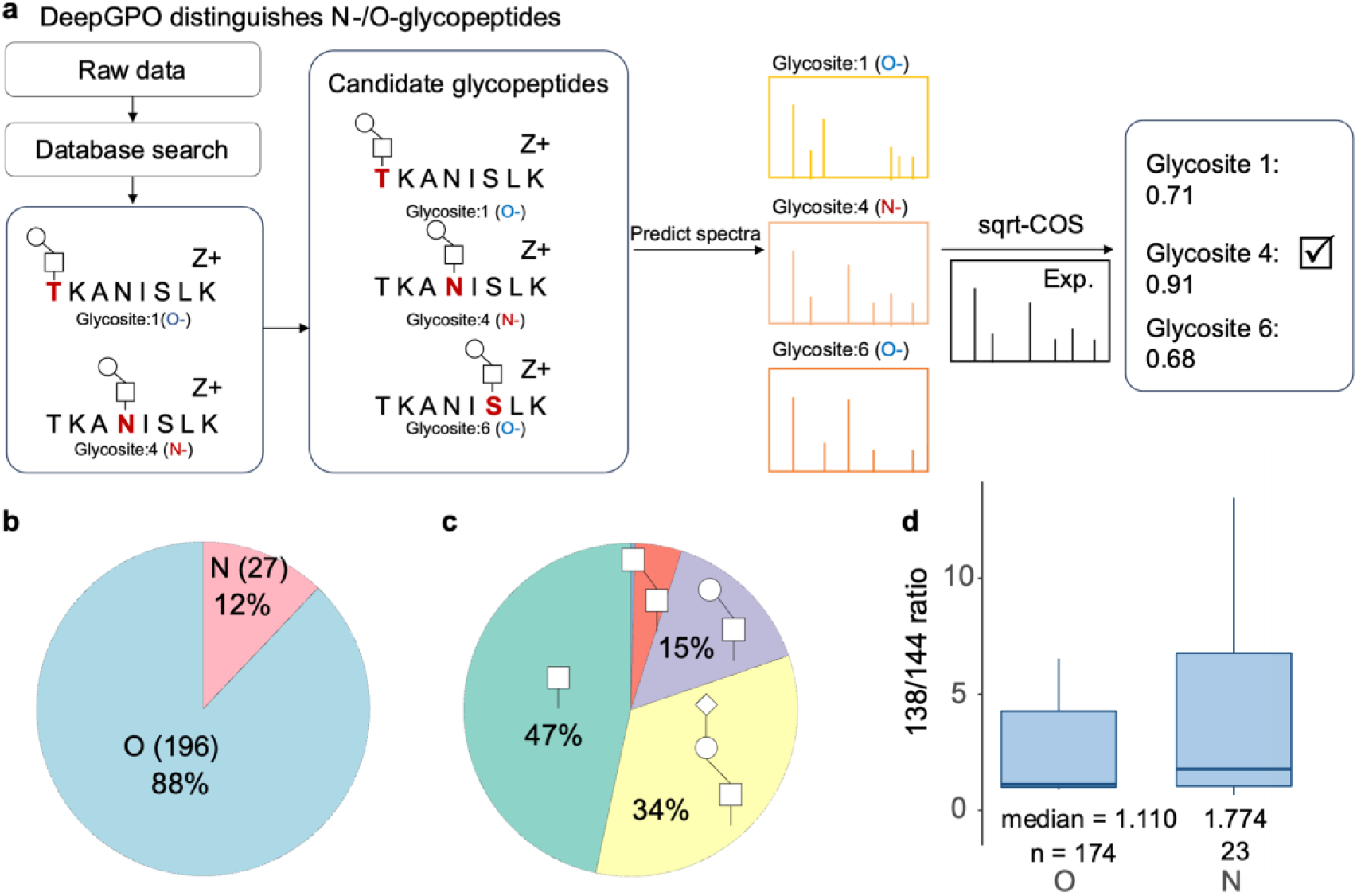
Performance of DeepGPO in *N*-/*O*-glycopeptide differentiation. (a) Protocol of performing *N*-/*O*-glycosite differentiation by DeepGPO. (b) Pie chart showing the number and proportion of identified *N*-glycopeptide PSMs and *O*-glycopeptide PSMs by DeepGPO. (c) Pie chart of the identified glycan types. (d) Intensity ratio of 138/144 peaks in the MS/MS spectra identified as *N*-glycopeptides and *O*-glycopeptides. Source data are provided as a Source Data file.

## Discussion

In this work, we present DeepGPO, a deep-learning model capable of accurately predicting MS/MS spectra for both *N*-glycopeptides and *O*-glycopeptides. Based on our previous DeepGP model for *N*-glycopeptides, DeepGPO adopts a number of advanced training techniques to mitigate the limited availability of *O*-glycopeptides training data. The introduction of a loss re-weighting method preserves a wider range of training MS/MS spectra while allowing the model to quantitatively learn from more reliable MS/MS spectra. Retaining all duplicated glycopeptide MS/MS spectra during model training ensures effective data augmentation, where the intrinsic variations in different MS/MS spectra for the same glycopeptide are captured by the deep-learning model. Since many fragmentation events in *N*-glycopeptides can also happen in *O*-glycopeptides, DeepGPO also largely benefits from the pre-trained parameters in DeepGP. Various test datasets were used in this study, including those with and without specific *O*-glycoprotease digestion, as well as data derived from both human and mouse samples. By strictly excluding glycopeptides available in the training data from the test data, high spectra similarity was still obtained between the experimental and predicted MS/MS spectra. It is noted that the current work focuses on the HCD MS/MS spectra prediction, as this fragmentation method is efficient and has been adopted by many studies.

With the successful prediction of *O*-glycopeptide MS/MS spectra, we assess the ability of localizing *O*-glycosylation by spectra matching. DeepGPO challenges the prevailing assumption that HCD fragmentation alone is insufficient for the localization of *O*-glycosites^27, 38^. Unlike traditional sequence searching, which relies on the observation of site-determine ions to identify *O*-glycosites, DeepGPO captures both m/z and intensity variations, offering superior performance in the analysis of *O*-glycopeptides. Three different datasets were used to demonstrate the principle, including one with IMPa *O*-glycoprotease digestion to generate *O*-glycopeptides starting with Ser/Thr and glycosylation at the N-termini, one with mucin-selective metalloprotease Amuc_0627 which cleaves between adjacent residues carrying truncated core 1 *O*-glycans, and one without any specific *O*-glycoprotease. Even without applying a strict similarity score threshold, DeepGPO yielded favorable results based solely on HCD spectra. When the similarity score threshold was set to 0.90 or 0.92, high consistency was observed between the DeepGPO result based on HCD and pGlyco3 result based on EThCD. The results demonstrate that DeepGPO together with HCD can effectively identify the *O*-glycosites, largely consistent with the enzyme specificity and the ETD based site-localization. With a similar concept, DeepGPO also demonstrates promising results in differentiating between *N*- and *O*-glycopeptides. Since HCD fragmentation can achieve greater coverage of glycopeptides due to faster cycle time and more efficient dissociation^27^, this advancement has the potential to enhance *O*-glycoproteomics.

The primary limitation of DeepGPO arises from the scarcity of data. Synthetic peptides are considered gold standards for studying fragmentation patterns, but there is a lack of datasets for synthetic *O*-glycopeptides. This scarcity hampers various applications, including the optimization of decoy generation methods and the calculation of false discovery rates (FDR) at both the glycan and glycopeptide levels. In our application of *O*-glycosites localization and differentiation of *N*- and *O*-glycopeptides, the lack of synthetic glycopeptides datasets also hampers further validation. Besides, the current work has primarily focused on singly glycosylated peptides, with preliminary investigations conducted on peptides bearing two glycan modifications. At present, the method is unable to effectively process peptides with three or more glycan attachments, owing to challenges such as limited training data and increased spectral complexity. With expanded datasets and the advancement of analytical tools, DeepGPO holds significant potential for broader application in glycoproteomics research.

In summary, DeepGPO is developed for the prediction of *N*-/*O*-glycopeptides MS/MS spectra, and has shown potential in localizing *O*-glycosites and differentiating *N*- and *O*-glycopeptides using HCD MS/MS spectra. With the development of glycoproteomics, particularly in mass spectrometry methods and data analysis software solutions, we anticipate that DeepGPO will enhance the understanding of the heterogeneity and complexity of glycoproteomics, and further aid biologists in the study of glycobiology.

## Methods

### Glycoproteome data analysis by pGlyco3

Glycoproteomic datasets were obtained from ProteomeXchange (https://www.proteomexchange.org/), and data analysis was performed using pGlyco3 (version: pGlyco3.1, https://github.com/pFindStudio/pGlyco3/releases). For *O*-glycoproteomics data analysis, HCD-pd-EThCD or EThCD spectra were analyzed using the pGlyco-*O-* Glycan.gdb glycan database; while for HCD spectra, the protein sequence FASTA file, glycan database, and modification settings were consistent with those described in the original publications of the corresponding datasets. For *N*-glycoproteomics data analysis, the glycan type was specified as *N*-glycan, and the glycan database was pGlyco-*N-*Human.gdb. Analytical parameters were set as follows: precursor mass tolerance at 5 ppm, fragment mass tolerance at 20 ppm, and glycopeptide FDR at 0.01. All other parameters were kept at their default settings.

### DeepGPO framework

The DeepGPO framework is based on DeepGP^23^. The performance of MS/MS spectra prediction is evaluated using sqrt-cosine similarity:

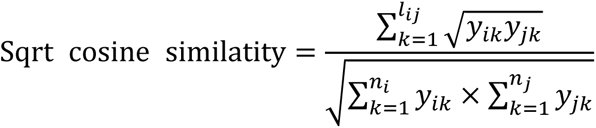

Here, *l_ij_* represents the number of common peaks between spectrum *i* and spectrum *j*, while *n_i_* and *n_j_* denote the counts of peaks identified as glycopeptide fragments in spectra *i* and *j*, respectively. The variable *y* indicates peak intensity. A tolerance of 20 ppm is applied to identify common peaks, and in cases where multiple peaks fall within this tolerance, the peak closest in distance is selected.

DeepGPO predicts 24 types of b/y fragment peaks, denoted as b1, b1n, b1o, b1h, b1nh, b1oh, y1, y1n, y1o, y1h, y1nh, y1oh, b2, b2n, b2o, b2h, b2nh, b2oh, y2, y2n, y2o, y2h, y2nh, y2oh. The first character indicates the ion type (b or y ions), followed by the fragment charge (1 or 2). The last character(s) specify the type of neutral loss: “o” represents the loss of H_2_O, “n” indicates loss of NH_3_, and “h” denotes loss of a monosaccharide HexNAc. In our analysis, peaks without the “h” symbol correspond to b/y fragments containing one HexNAc moiety, while those with the “h” symbol represent b/y fragments that have lost the HexNAc moiety. This nomenclature is designed to ensure that each symbol clearly indicates the type of loss, simplifying coding.

DeepGPO recognizes 16 types of B/Y fragment peaks (B1, B1n, B1o, B1f, Y1, Y1n, Y1o, Y1f, B2, B2n, B2o, B2f, Y2, Y2n, Y2o, Y2f). The first character indicates the ion type (B or Y ions), and "f" signifies the loss of a monosaccharide Fuc, with the other symbols adhering to the same nomenclature rules of b/y fragment peaks. The total number of peaks is determined by the number of cleavage events and the types of ions associated with each event. Notably, ions that are theoretically impossible are not excluded from model predictions; instead, predicting these ions results in penalties within our scoring system, which encourages the model to refine its learning. However, during the post-processing phase, ions identified as theoretically impossible are omitted when converting the output matrix into the corresponding predicted MS/MS spectra. Importantly, there are no restrictions on precursor charge states, but we consider fragment charges up to +2.

### DeepGPO model training

DeepGPO uses the search results by pGlyco3 for model training, validation and test. Glycopeptides present in the training data are excluded from the validation and test data for MS/MS prediction performance evaluation. The weight of the MS/MS spectra used for model training should exceed 0.5 (**Supplementary Fig. 2**). During model training, the MS/MS spectra intensity is normalized by the highest peak in the spectrum. The model utilizes a modified Mean Squared Error (MSE) loss function. Specifically, for spectra with assigned weights, the loss function is multiplied by the corresponding weight. Training of DeepGPO on Dataset 1 took 1 hour using a single RTX 3090 GPU with 100 epochs, a batch size of 256, and a learning rate of 1×10^−4^. DeepGPO took 1 seconds to predict 300 spectra using a single RTX 3090 GPU.

### DeepGPO evaluation

The experimental MS/MS spectra are converted to Mascot generic format (.mgf) from the searching process of pGlyco3. DeepGPO generates the predicted MS/MS spectra. A tolerance of 20 ppm is established to identify common peaks between the experimental and predicted MS/MS spectra. When multiple peaks fall within this range, the peak with the shortest distance is selected. Any discrepancies between the compared spectra result in the missing peak intensity being assigned a value of 0, followed by the calculation of spectrum similarity. DeepGPO provides various metrics for this assessment, including square root cosine similarity (sqrt-cosine), cosine similarity (COS), and the pearson correlation coefficient (PCC).

### Analysis of doubly glycosylated peptides

For the analysis of doubly glycosylated peptides, the identification results were obtained from the original study using Byonic. The same filtering criteria as in the original study were applied to ensure consistency and comparability. Specifically, peptide-spectrum matches (PSMs) were retained if they met the following conditions: a Byonic score ≥ 200, a logProb value ≥ 2, and a peptide length of more than four amino acid residues.

The model framework of DeepGPO for doubly glycosylated peptides is consistent with that used for singly glycosylated peptides. A minor difference is that DeepGPO predicts 36 types of b/y fragment ions, including: b1, b1n, b1o, b1h, b1nh, b1oh, b1hh, b1nhh, b1ohh, y1, y1n, y1o, y1h, y1nh, y1oh, y1hh, y1nhh, y1ohh, b2, b2n, b2o, b2h, b2nh, b2oh, b2hh, b2nhh, b2ohh, y2, y2n, y2o, y2h, y2nh, y2oh, y2hh, y2nhh, y2ohh. The interpretation of the characters remains the same as in the singly glycosylated peptide model.

### Implementation and visualization

DeepGPO was developed using python (3.8.3, Anaconda distribution version 5.3.1, https://www.anaconda.com/) with the following packages: bidict (0.22.0), dgl (2.4.0+cu118), FastNLP (0.6.0), numpy (1.18.5), pandas (1.0.5), pyteomics (4.7.3), pytorch (1.8.1), torchinfo (1.7.1), transformers (4.12.5), scikit-learn (1.5.2) and scipy (1.14.1). Visualization was performed using custom scripts in R (4.0.2) with the following packages: VennDiagram (1.6.20) and ggplot2 (3.3.2). The mirrored spectra are plotted by GP-plotter^39^ (1.0.0).

## Data availability

The raw datasets used in this study are available in the PRIDE^40^ database under accession code PXD037415^27^, PXD018560^28^, PXD004590^29^, PXD032164^34^, PXD022896^30^, PXD009476^31^, PXD020077^9^, PXD039583^32^, PXD027616^33^, PXD031225^34^, PXD035775^35^, PXD017646^36^, and PXD024995^38^. Further details regarding the datasets and raw files used in this study can be found in **Supplementary Table 1** and **Supplementary Note 1**. The source data underlying all figures except for those not including statistics are provided as a Source Data file.

## Code availability

DeepGPO along with the user guide is freely available via https://gitfront.io/r/yuz2011/Ah4BPe9TJ4bG/DeepGPO/ and GitHub [https://github.com/lmsac/DeepGPO] (*The GitHub page is currently private, and will be released upon publication of the manuscript.*). A user-friendly software solution, along with a demonstration video, example inputs and pre-trained models are available: https://drive.google.com/drive/folders/1frGwRgUSPbBMk27uYDrf4wdqn6KSzEjb

## Acknowledgements

This work was supported by the Science and Technology Commission of Shanghai Municipality (23JS1400100), and the National Natural Science Foundation of China (NSFC 22374031).

## Author contributions

Y.Z. did the majority of coding work and data analysis, and wrote the original draft of the manuscript. Y.W. built the deep-learning model. L.Q. supervised all aspects of the work, and finalized the manuscript. All authors were involved in the design of this work.

## Supplementary Information

**Supplementary Table**, **Figures** and **Note** are in the supplementary PDF file.

**Supplementary Data 1**: Identification of O-glycopeptides by DeepGPO and pGlyco3 from the test data of Dataset 1 at 5% empirical FLR.

**Supplementary Data 2**: Identification of *O*-glycopeptides by DeepGPO from Dataset 5.

**Supplementary Data 3**: The result of DeepGPO for *N*-/*O*-glycosite differentiation for Dataset 13.

