## Supplementary Information for "Deep Learning Prediction of Intact *N*-/*O*-Glycopeptide Tandem Mass Spectra Enhances Glycoproteomics"

Zong et al.

**Supplementary Table 1.** Datasets for deep learning model training, evaluation and test.

| Name | Instrument | Sample type | *O*-glycoproteases | Accession |
|---|---|---|---|---|
| Dataset 1 | Orbitrap Fusion Lumos Tribrid | Human / Mouse | IMPa/None | PXD037415[1] |
| Dataset 2 | LTQ-Orbitrap Velos / Orbitrap Fusion Tribrid | Human / Pig | None | PXD018560[2] |
| Dataset 3 | Orbitrap Fusion Tribrid | Human | None | PXD004590[3] |
| Dataset 4 | Orbitrap Fusion Tribrid | Mucin | AM0627 | PXD032164[4] |
| Dataset 5 | Orbitrap Fusion Lumos Tribrid | SARS-CoV-2 | None | PXD022896[5] |
| Dataset 6 | Q-Exactive HF | Human | OgpA | PXD009476[6] |
| Dataset 7 | Orbitrap Fusion Tribrid | Bovine / Human | OgpA | PXD020077[7] |
| Dataset 8 | Orbitrap Eclipse Tribrid | T cell immunoglobulin and mucin-domain-containing (TIM) proteins | IMPa/OgpA/SmE/None | PXD039583[8] |
| Dataset 9 | Orbitrap Fusion Tribrid | Mucin | None | PXD027616[9] |
| Dataset 10 | Orbitrap Fusion Tribrid | Mucin | AM0627 | PXD031225[4] |
| Dataset 11 | Orbitrap Fusion Tribrid | Mucin | IMPa/OgpA/StcE | PXD035775[10] |
| Dataset 12 | Orbitrap Fusion | Mucin | StcE | PXD017646[11] |
| Dataset 13 | Orbitrap Fusion Tribrid | Human | None | PXD024995[12] |

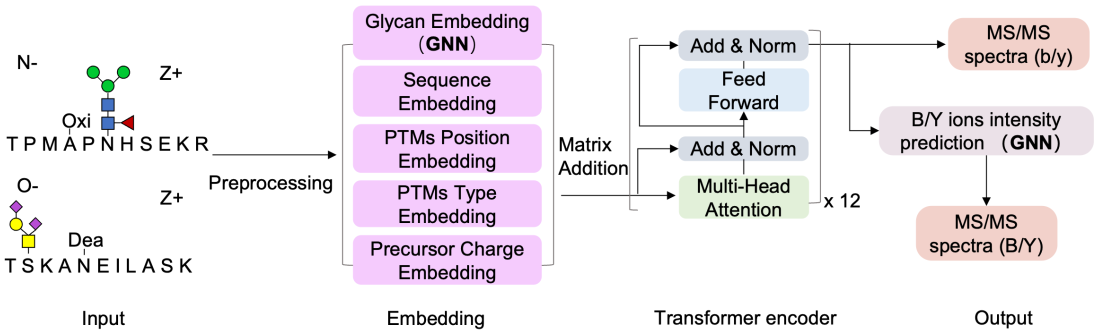


**Supplementary Figure 1.** The model architecture of DeepGPO.

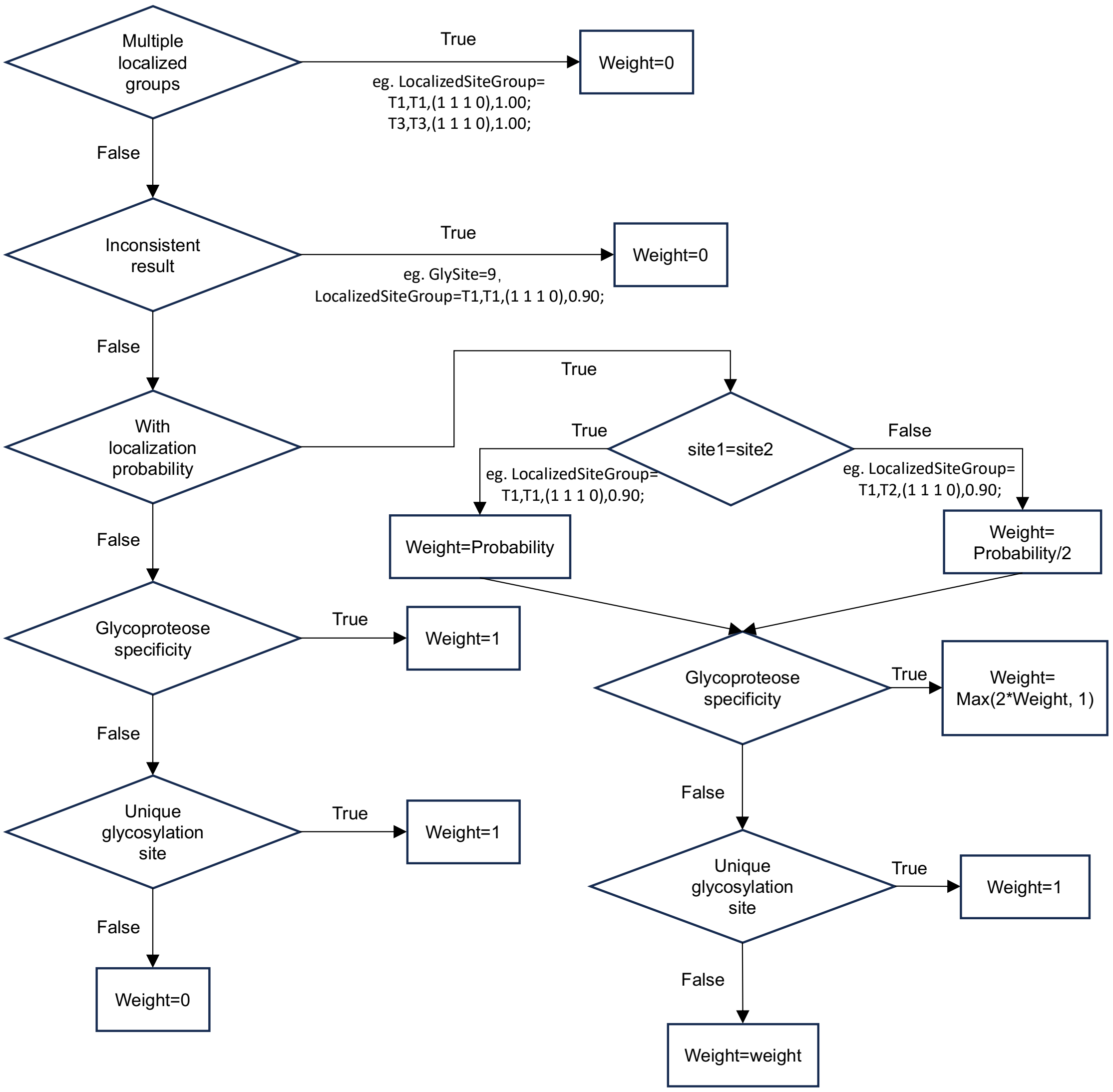


**Supplementary Figure 2.** Workflow for assigning training weights to glycopeptide MS/MS spectra. The weights are only applied to the mono-glycosylated peptides. An initial weight is obtained from the pGlyco3 as the localization probability. The MS/MS spectra identified as multi-glycosylated peptides receive the weight of 0. When the reported glycosite is inconsistent with those in the LocalizedSiteGroup, the weight is 0. Spectra without localization probability due to the lack of ETD data are assigned the weight of 0. If site1 is the same as site2 in the LocalizedSiteGroup, the weight is the probability. If site1 differs from site2, the weight of each site is halved. When specific glycoprotease is used, for pGlyco3 identification results that match glycoprotease specificity, the weight is double of the weight without considering the glycoprotease specificity, ensuring a minimum weight of 1. Glycopeptides with only one potential glycosylation site receive the weight of 1.

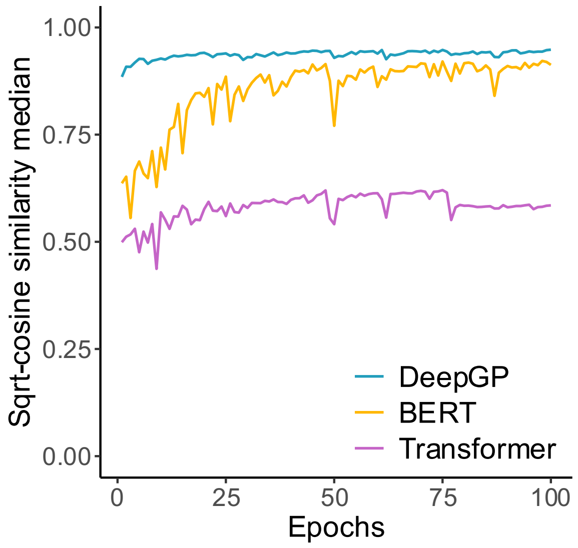


**Supplementary Figure 3.** Performance of DeepGP, BERT and Transformer as the base models for DeepGPO training using Dataset 1. Source data are provided as a Source Data file.

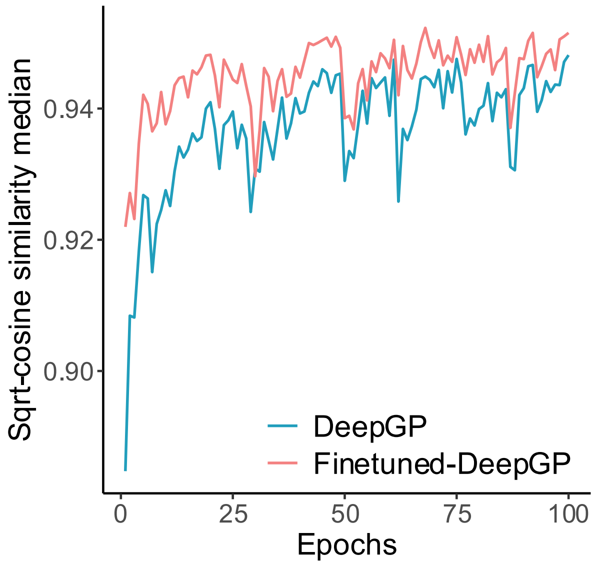


**Supplementary Figure 4.** Performance of DeepGP and Finetuned-DeepGP as the base models for DeepGPO training using Dataset 1. DeepGP means that the base model is DeepGP and Finetuned-DeepGP means that the base model is DeepGP trained with other *O*-glycopeptides datasets (Dataset 2-11). Source data are provided as a Source Data file.

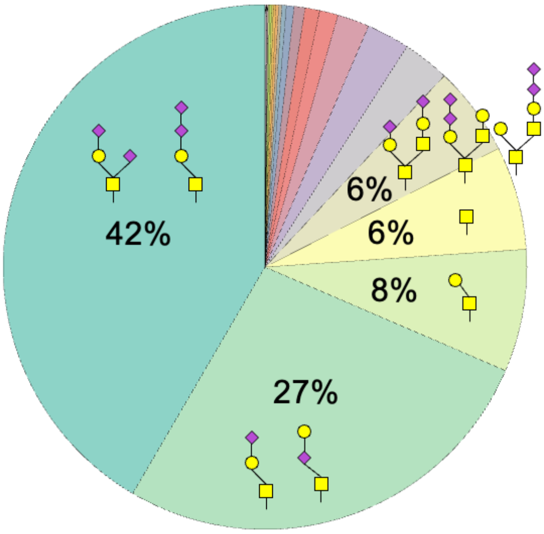


**Supplementary Figure 5.** The glycan types for the glycopeptides identified from the test data of Dataset 1.

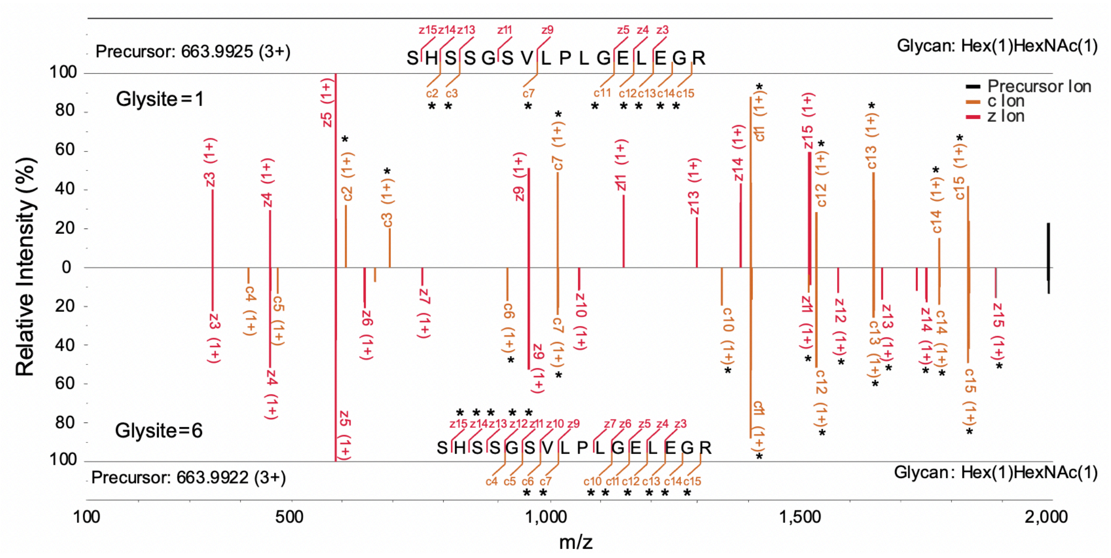


**Supplementary Figure 6.** Comparison of the experimental ETD MS/MS of the glycopeptides of the same peptide sequence and glycan composition but different glycosites. Top: the experimental MS/MS spectra with the glycosite at the first position; Bottom: the experimental MS/MS spectra with the glycosite at the sixth position. *: fragments with the intact glycan. Experimental MS/MS spectra were pre-processed to remove noise peaks for clearer comparison.

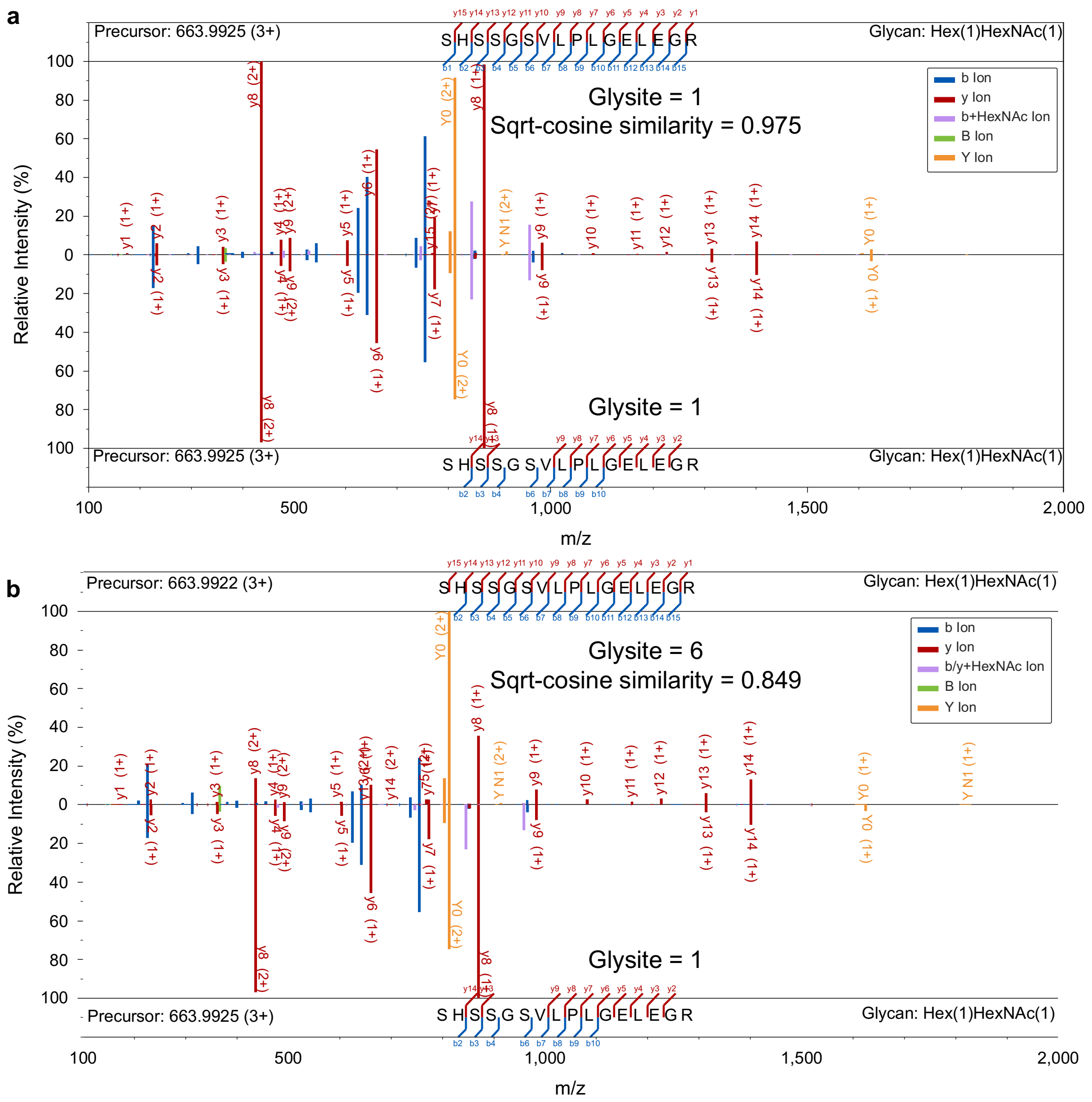


**Supplementary Figure 7.** Comparison of the experimental HCD MS/MS with the predicted HCD MS/MS for the glycopeptides of the same peptide sequence and glycan composition but either glycosite at the first position or the sixth position. (a) Glycosite = 1 for both predicted and experimental MS/MS spectra; (b) Glycosite = 6 for predicted MS/MS spectra and Glycosite = 1 for experimental MS/MS spectra. Top: the predicted MS/MS spectra; Bottom: the experimental MS/MS spectra. Experimental spectra were pre-processed to remove noise peaks for clearer comparison.

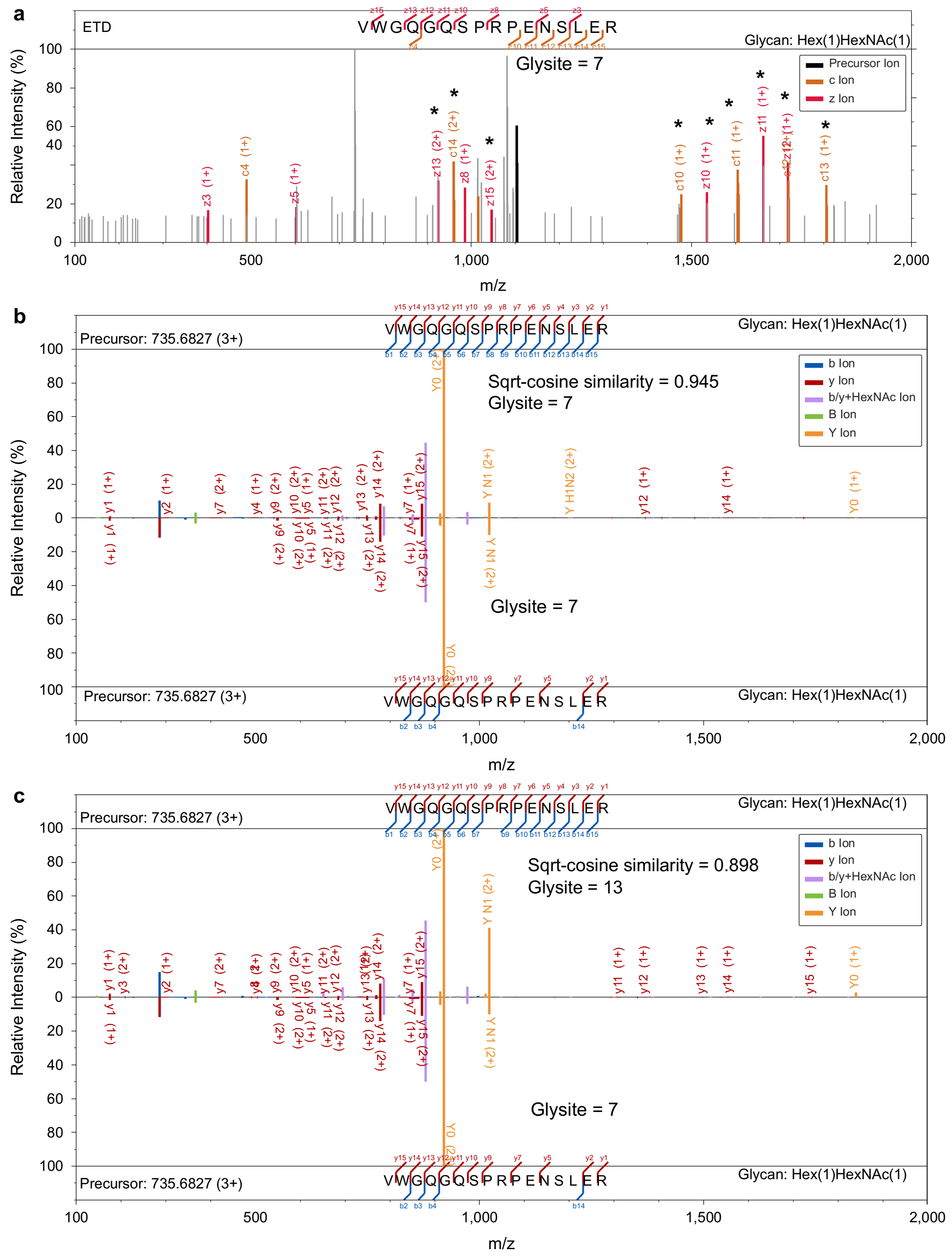


**Supplementary Figure 8.** (a) The ETD MS/MS spectrum of a glycopeptide, demonstrating the glycosylation at site 7. (b) Comparison between the predicted HCD MS/MS spectrum (top) of the glycopeptide with site at 7 to the experimental HCD MS/MS spectrum (bottom) of the glycopeptide. (c) Comparison between the predicted HCD MS/MS spectrum (top) of a glycopeptide with the same peptide sequence and glycan composition but a site at 13 to the experimental HCD MS/MS spectrum (bottom) of the glycopeptide. Experimental MS/MS spectra were pre-processed to remove noise peaks for clearer comparison.

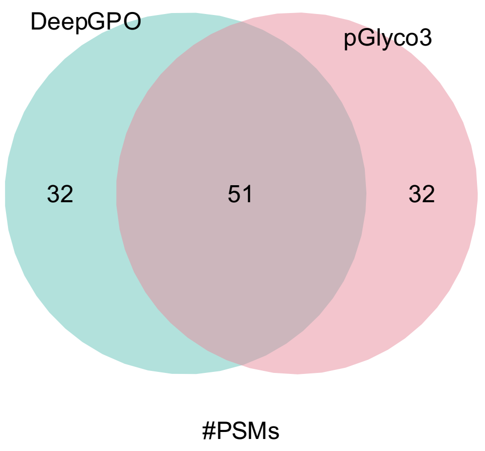


**Supplementary Figure 9.** Venn diagram of the number of PSMs identified by DeepGPO without cutoff of sqrt cosine similarity and by pGlyco3 with glycosite localization probability < 0.8 for Dataset 3.

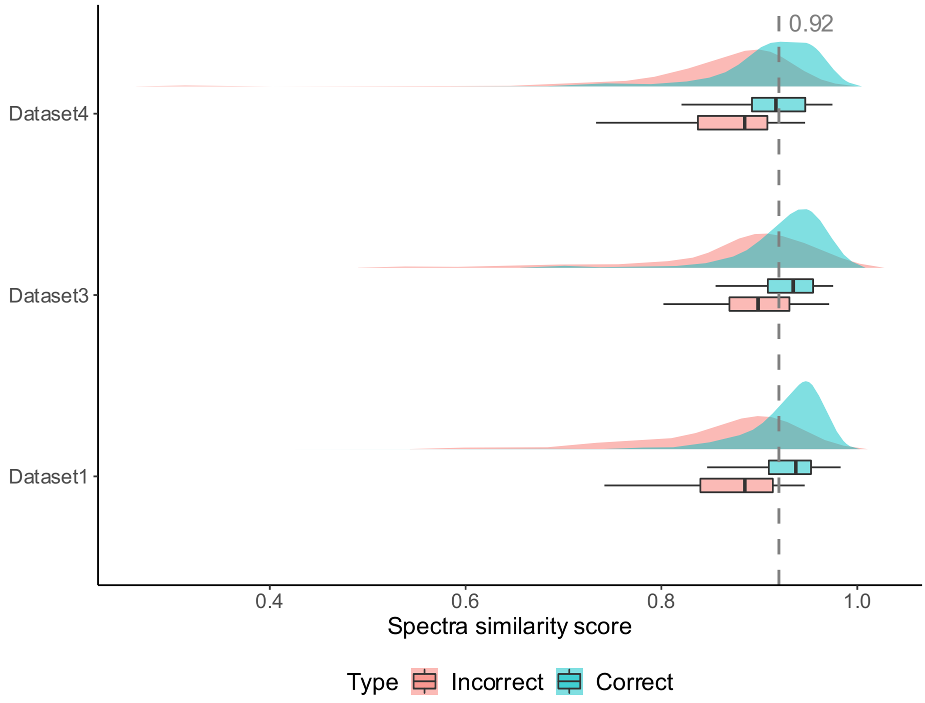


**Supplementary Figure 10. The score distributions (sqrt cosine similarity) for both correct and incorrect PSMs from three datasets (Dataset 1, Dataset 3 and Dataset 4).** Half violin graph (Top) and boxplot (Bottom) of sqrt cosine similarity distribution for correct and incorrect glycopeptides hits. Boxes mark the first and third quartile, with the median highlighted as the line, and whiskers mark the minimum/maximum values within the 1.5 interquartile range. Outliers are not shown. Source data are provided as a Source Data file.

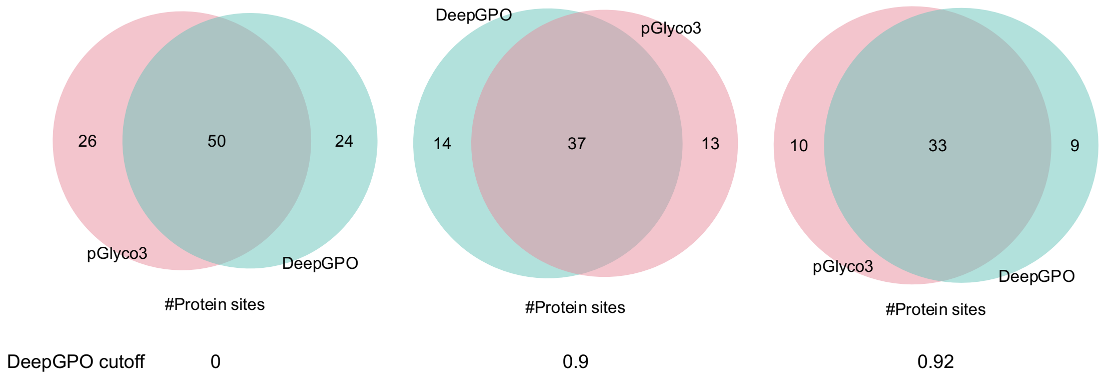


**Supplementary Figure 11.** Venn diagram of the number of protein glycosites identified by DeepGPO with a similarity score cutoff of 0, 0.9 and 0.92, and by pGlyco3 for Dataset 3. The pGlyco3 glycosite localization probability was ≥ 0.8. pGlyco3 analyzed the EThcD while the DeepGPO analyzed the HCD spectra from the EThcD.

**Supplementary Note 1: Description of the datasets used.**

DeepGPO was mainly benchmarked on Dataset 1, Dataset 2, Dataset 3, Dataset 4, Dataset 5, Dataset 12, and Dataset 13. Dataset 1 is accessed through ProteomeXchange with identifier PXD037415 previously published by Suttapitugsakul et al.[1] *O*-glycopeptides from HEK293 cells or mouse brain tissues were released by treating the samples with IMPa *O*-glycoprotease. For the HEK293 cells, HCD-pd-EThCD or HCD alone was used for glycopeptide fragmentation. For the mouse brain tissues, only HCD-pd-EThCD was used for glycopeptide fragmentation and there were (a) single-shot experiments, (b) experiments where the peptide load was increased, and (c) experiments with a modified MS1 scan range. The same mouse brain tissue was also used for both *N*- and *O*-glycoproteomics analyses including (a) *O*-glycoproteomics analyses using HCD-pd-EthCD, and (b) *N*-glycoproteomics analyses using sceHCD with collision energies of 20%, 30% and 40%.

Dataset 2 is accessed through ProteomeXchange with identifier PXD018560 previously published by Madsen et al.[2] In this study, the authors of the original publication used samples from human, rat and pig. In our study, datasets of human and pig were used. ETD triggering of subsequent HCD scan was used for the experiments.

Dataset 3 is accessed through ProteomeXchange with identifier PXD004590 previously published by King et al.[3] *O*-glycopeptides from human plasma, platelets and endothelial cells were treated with trypsin and/or chymotrypsin. MS/MS analysis was performed using HCD and ETD.

Dataset 4 is accessed through ProteomeXchange with identifier PXD032164 previously published by Shon et al.[4] Digestion was performed with AM0627 or AM0627 mutants (AM0627$^{W149A}$, AM0627$^{F290A}$, AM0627$^{Y287A}$) against recombinant glycoproteins, podocalyxin, MUC16, PSGL-1, and CD43. The glycopeptides were analyzed by HCD-pd-EThCD.

Dataset 5 is accessed through ProteomeXchange with identifier PXD022896 previously published by Zhang et al.[5] *O*-glycopeptides were from SARS-CoV-2 S proteins treated with trypsin and trypsin/Glu-C after de-*N*-glycosylation using PNGase F. All the samples were analyzed using HCD and EthCD.

Dataset 12 is accessed through ProteomeXchange with identifier PXD017646 previously published by Riley et al.[11]. In this study, multiple dissociation methods were employed for glycoproteomics analysis. We analyzed the *O*-glycopeptide data generated using HCD-pd-ETD and HCD-pd-EThcD dissociation methods. For our investigation of *O*-glycopeptides, we utilized the Byonic search results provided in the original publication.

Dataset 13 is accessed through ProteomeXchange with identifier PXD024995 previously published by Malaker et al.[12]. *O*-glycopeptides of mucins were enriched from complex samples like cell lysate and crude ovarian cancer patient ascites fluid for LC-MS/MS analysis using HCD.
